# Age- and sex- dependent effects of moderate gestational day 12 prenatal alcohol exposure on anxiety-like behaviors, ethanol intake, and mechanical sensitivity

**DOI:** 10.64898/2026.05.19.726255

**Authors:** Sarah E. Winchester, Elena I Varlinskaya, Marvin R. Diaz

**Author notes:** **Address correspondence to:** Dr. Marvin R. Diaz, Department of Psychology, Binghamton University PO Box 6000, State University of New York, Binghamton, NY 13902-6000.

## Abstract

**Rationale:** Prenatal alcohol exposure (PAE) can result in Fetal Alcohol Spectrum Disorder (FASD), which consists of a group of diagnosable medical conditions that can include an increased risk for anxiety disorders and/or alcohol misuse, and sensory issues, such as increased mechanical sensitivity.

**Objective:** This study investigated how a single moderate PAE on gestational day 12 (G12) alters anxiety-like behavior, ethanol (EtOH) intake, and mechanical sensitivity across the lifespan of Sprague Dawley rats.

**Methods:** Pregnant dams were exposed to vaporized EtOH or room air (control) for 6 hours (BECs ∼108 mg/dL). Testing in male and female offspring began at three different ages: juveniles (∼postnatal day (P) 25), adolescents (∼P45) and adults (∼P80).

**Results:** The greatest PAE effects were observed in adolescent animals, with alterations in anxiety-like behaviors demonstrated in the light-dark box and elevated plus maze. Additionally, adolescent female animals consumed more sweetened EtOH compared to males. However, PAE adolescent animals consuming less sweetened EtOH compared to their counterparts, which was also observed in adult PAE females. Interestingly, this effect is reversed in juvenile and adolescent males when tested with unsweetened EtOH, with juvenile females consuming more EtOH also. Finally, PAE and air animals exhibited increased mechanical sensitivity following post-natal EtOH consumption across all ages.

**Conclusion:** These data demonstrate that there are age- and sex-specific effects of PAE on anxiety-like behaviors, EtOH intake, and mechanical sensitivity that are more distinct in adolescent animals.

## Introduction

Alcohol consumption by pregnant women in the U.S. continues to increase; with a 2% increase from previous years, in 2018-2020 the CDC reported that 1 in 7 (13.5%) pregnant women consumed alcohol in the last 30 days, while 1 in 20 (5.2%) engaged in binge drinking (Gosdin et al., 2022). Alcohol consumed by a pregnant woman passes through the placental barrier, and the fetus’ blood alcohol level can match maternal levels within 2 hours of consumption, suggesting that fetal alcohol levels are sustained longer due to slower metabolism (Dejong et al., 2019). This teratogenic exposure can result in the development of Fetal Alcohol Spectrum Disorder (FASD), which presents as a wide range of long-lasting adverse effects.

One of the most debilitating consequences of prenatal alcohol exposure (PAE) is the higher prevalence for developing an anxiety disorder, affecting approximately 20% of those with FASD, with anxiety symptoms presenting in early childhood and persisting throughout adulthood (Leech et al., 2006; O’Connor and Paley, 2009; Steinhausen et al., 2003). Additionally, exposure to alcohol *in utero* predicts future alcohol misuse, with onset typically occurring in adolescence and lasting throughout adulthood (Baer et al., 1998; Baer et al., 2003). Misuse of alcohol is positively correlated with anxiety, as individuals diagnosed with an anxiety disorder are 2.3 times more likely to misuse alcohol (Anker and Kushner, 2019). The relationship between alcohol misuse and anxiety is bidirectional, with each capable of exacerbating the other, creating a vicious comorbid cycle.

Interestingly, misuse of alcohol also intricately interacts with heightened pain sensitivity. Individuals who misuse alcohol are 43% more likely to experience moderate-to-severe pain (Brennan and Soohoo, 2013; McDermott et al., 2018). Coincidentally, psychiatric disorders, like anxiety, are highly comorbid in patients with pain, as persistent pain may contribute to anxiety. In support of this, individuals suffering from persistent pain are four times more likely to be diagnosed with an anxiety disorder (Ditre et al., 2019). Clearly, there are reciprocal relationships among alcohol use, anxiety, and pain, in which each condition exacerbates the others.

Importantly, research has not examined how PAE may alter the relationships between anxiety, alcohol consumption, and pain. To date, research has shown that individuals with PAE exhibit abnormal sensory-processing, such as exacerbated sensitivity to light touch, termed allodynia (Franklin et al., 2008). Pre-clinical research using a *moderate* PAE throughout gestation [blood ethanol concentrations (BECs) in dams ∼60-80 mg/dL] has found that allodynia was exacerbated after mild sciatic chronic constriction injury only in subjects with PAE, which has been associated with alterations in neuroimmune function (Noor and Milligan, 2018; Noor et al., 2017; Pasmay et al., 2025; Pritha et al., 2026; Sanchez et al., 2017). This suggests that PAE individuals may have a hyper-reactive response after a second “hit” to their nervous system, perhaps due to the teratogenic effects of alcohol during development of the central nervous system.

Animal models have demonstrated that altered developmental trajectories observed following PAE are dependent on the level and timing of exposure to alcohol. In rodents, gestational day 12 (G12; equivalent to 2^nd^ trimester in humans) is a crucial time period for the development of the amygdala, a brain structure involved in anxiety-like behaviors, alcohol-related behaviors, and pain processing (Allen et al., 2021; See review: Gilpin et al., 2015; Soma et al., 2009). In support of the relevance of G12, our lab has shown that *adult* male and female offspring exposed to a heavy PAE (BECs >300 mg/dL) on G12 exhibit increased ethanol (EtOH) intake regardless of context (social or alone drinking) (Diaz et al., 2020). Given that moderate PAE is more common in humans (Ethen et al., 2009; Gosdin et al., 2022), our lab has also found that *adolescent* males (P40-48) with a history of G12 moderate PAE (BECs ∼76-90 mg/dL) exhibit enhanced anxiety-like behavior in a number of assays measuring anxiety, including novelty induced hypophagia (NIH), light dark box (LDB), and open field (OF), with no effects of moderate PAE evident in *adolescent* females (Rouzer et al., 2017). Furthermore, the same study found that PAE *adult* males buried significantly less marbles, suggesting that anxiety-like behaviors had ameliorated and were lower than in controls by adulthood (Rouzer et al., 2017), consistent with reduced anxiety-like behavior in the elevate plus maze observed in heavy G12 PAE adult offspring (Diaz et al., 2016).

Taken together, these findings highlight the importance of exploring PAE’s influence on nonsocial anxiety, EtOH intake, and pain sensitivity. Furthermore, preclinical research has shown that effects of PAE on emotional processing, EtOH acceptance, and, perhaps, pain sensitivity are sex- and age-dependent. Therefore, the present study was designed to assess the effects of G12 moderate PAE on anxiety-like behavior, subsequent EtOH intake, and mechanical sensitivity across ontogeny in male and female offspring.

## Materials and Methods

### Subjects

Experimental subjects were the offspring of Sprague-Dawley rats that were bred in our colony at Binghamton University from breeders purchased from Envigo/Inotiv (Indianapolis, IN, USA). For breeding, after a 2-week acclimation period, two adult females and one male were housed together for four days, and females were checked daily for sperm via vaginal smears, with the first day of detectable sperm designated as G1 as described (Mooney and Varlinskaya, 2011; Rouzer et al., 2017; Rouzer and Diaz, 2021, 2022; Winchester and Diaz, 2025). Pregnant dams were single-housed and weighed throughout gestation (G1, G10, G20). All litters were culled to 12 pups, maintaining a 1:1 male to female ratio, when possible. Pups were weighed on post-natal day (P) 2, P7 and P12. On P21, pups were weaned and pair-housed with a same-sex litter mate. A total of 67 litters was used (air = 31; PAE = 30; naïve = 3). All animals were held in a temperature-controlled colony (22ℒC), on a 12h light-dark cycle (lights on from 0700 to 1900) with *ad libitum* access to food (non-breeders: Purina 5LOD rat chow laboratory diet; breeders: Purina 5008 rat chow laboratory diet) and water. All experimental procedures and maintenance of rats were in accordance with the guidelines for animal care established by the National Institutes of Health, and all experimental protocols were approved by Binghamton University’s Institutional Animal Care and Use Committee (IACUC).

### Moderate prenatal alcohol exposure (mPAE)

Pregnant females were transferred to novel cages and relocated to vapor chambers on G12, as previously described (Przybysz et al., 2023; Rouzer et al., 2017; Rouzer and Diaz, 2022; Winchester and Diaz, 2025). Pregnant females were exposed to vaporized EtOH (95%) or room air (control) for 6 h, with this exposure resulting in a peak BEC of ∼108 mg/dL (Winchester and Diaz, 2025). Animals had *ad libitum* access to food and water throughout the duration of the exposure. Upon completion of the exposure, females were transferred to a clean cage with new food and water to prevent further EtOH exposure. Females were left undisturbed, apart from weighing, until parturition.

### Light Dark Box (LDB)

LDB testing was used to assess anxiety-like behaviors. This apparatus consisted of two attached compartments (34 x 24 x 24 cm) joined by a circular opening in the middle where the animal could easily access either compartment as previously described (Rouzer et al., 2017). The light compartment of this apparatus was made of white plexiglass with a clear lid that allowed room light in (∼90 lux; digital lux meter: Dr. Meter, Model no. LX1330B). The dark compartment was made of black plexiglass with a black lid that allowed no light in. The apparatus was cleaned with 3% hydrogen peroxide between each trial. At the beginning of testing, animals were placed in the light compartment facing away from the circular opening. Animals were video recorded and allowed to explore the apparatus for 5 minutes. Recordings were scored for: time spent in light compartment, egress latency (time it took to enter dark compartment for the first time), and head pokes. An entry into a compartment was recorded when an experimental subject placed all four paws in that compartment, and head pokes from the dark compartment to the light compartment were measured when both ears passed through the circular opening.

### Elevated Plus Maze (EPM)

EPM was another test used to assess anxiety-like behaviors. Juveniles and adolescents were tested in an apparatus measuring: length of arms (36.5 cm), width of arms (9.7 cm), wall height (24.1 cm), and plastic edges along the open arms (0.63 cm). Adults were tested in an apparatus measuring: length of arms (48.3 cm), width of arms (12.7 cm), and wall height (29.2 cm) as previously described (Rouzer et al., 2017). Open arms of the apparatus were illuminated at ∼90 lux. Pretesting social isolation has been shown to promote exploration (Doremus-Fitzwater et al., 2009); therefore, immediately before testing, animals were isolated in a novel cage for one hour. Following this, experimental subjects were placed in the center of the maze facing an open arm. The 5-minute EPM test was video recorded for later scoring. The apparatus was cleaned with 3% hydrogen peroxide between each trial. Behavioral measures scored and analyzed included percent of time spent in the open arms, percent open arm entries, closed arm entries, head dips, and stretch attend postures (Doremus-Fitzwater et al., 2009; Przybysz et al., 2020).

### Ethanol (EtOH) Intake

The day after behavioral testing, animals were given a 30-minute limited access to EtOH on Monday, Wednesday, and Friday for 2 cycles as previously described (Diaz et al., 2020). In the *first* EtOH experiment, animals were transported to behavior rooms where they were placed alone in holding cages and given access to 10% EtOH in super-sac (SS) solution (3% sucrose + 0.125% saccharin). EtOH + SS solution was made fresh before every testing session. In the *second* EtOH experiment, a different subset of animals was given access to 10% EtOH, with no SS solution. EtOH bottles were labeled with animals’ ID number and filled with ∼ 40 mL of solution. Each day, each bottle was weighed [grams (g)] before testing began, and each animal’s weight (g) was taken. After 30 minutes of access, bottles were weighed (g) again and animals were placed back in home cages with cage mates and returned to the colony.

### Supersac (SS) Intake

In another subset of animals, an additional assessment of SS intake was used to determine whether consumption of EtOH + SS was influenced by the sweeteners used in the solution (sucrose and saccharin). We tested SS intake and EtOH intake separately, allowing us to compare their consumption patterns, even though the two solutions were not presented simultaneously. Animals were exposed to SS solution using the same 30-minute limited access paradigm described above.

### Von Frey Test

Exacta touch test ® Von Frey filaments were used to assess thresholds for mechanical allodynia. These are a set of 20 monofilaments with a range target force indicated in grams (g): 0.008 g, 0.02 g, 0.04 g, 0.07 g, 0.16 g, 0.4 g, 0.6 g, 1 g, 2 g, 4 g, 6 g, 8 g, 10 g, 15 g, 26 g, 60 g, 100 g, 180 g, and 300 g. Animals were transferred from the colony room to a dimly lit room where they were habituated for 2-consecutive days (30 minute session/day) before testing. Animals were placed on top of a metal wire grid (36.8 x 50.8 cm; inter-bar space: 2.2 cm) elevated by plexiglass containers for access to hind paws (20.3 cm). A plexiglass container (47.3 x 25.4 x 20.3 cm) was placed over the apparatus to contain the animal. Monofilaments were applied to the plantar surface of the left or right hind paw. Left and right paws were randomized between animals, and the same paw was used for each testing session. Baseline assessment was initiated with the 2 g monofilament; each filament was applied 3 times to test for a response. A response was considered a paw withdrawal, shaking of the paw, or licking of the paw. If there was no response to a filament, then the next larger filament would be presented to the hind paw, if there was a response then the next descending filament would be presented. To record a response, an animal needed to respond to a filament three times consecutively. All equipment was cleaned with 3% hydrogen peroxide between animals and at the completion of testing day.

### Blood Ethanol Concentrations (BECs)

To ensure accurate measurements of EtOH intake in the initial drinking experiment (limited-access drinking paradigm with EtOH *+* SS), tail blood collection was performed at the conclusion of the 6th drinking session. All blood samples were analyzed using an Analox AM1 Alcohol Analyser (Analox Instruments Ltd, The Vale, London).

### Experimental Design (Experimental timeline: Figure 1)

To avoid litter effects, no more than 2 animals per sex per litter were randomly assigned to an age group. All behavior experiments were performed in different subsets of juveniles (∼P25-30), late-adolescents (∼P45-50), and adults (∼P80+). Specifically, each age group had its own set of subjects that were used separately, so no group was tested more than once across the different experiments. All testing began within the described age ranges and continued for 2 weeks.

**Figure 1:**
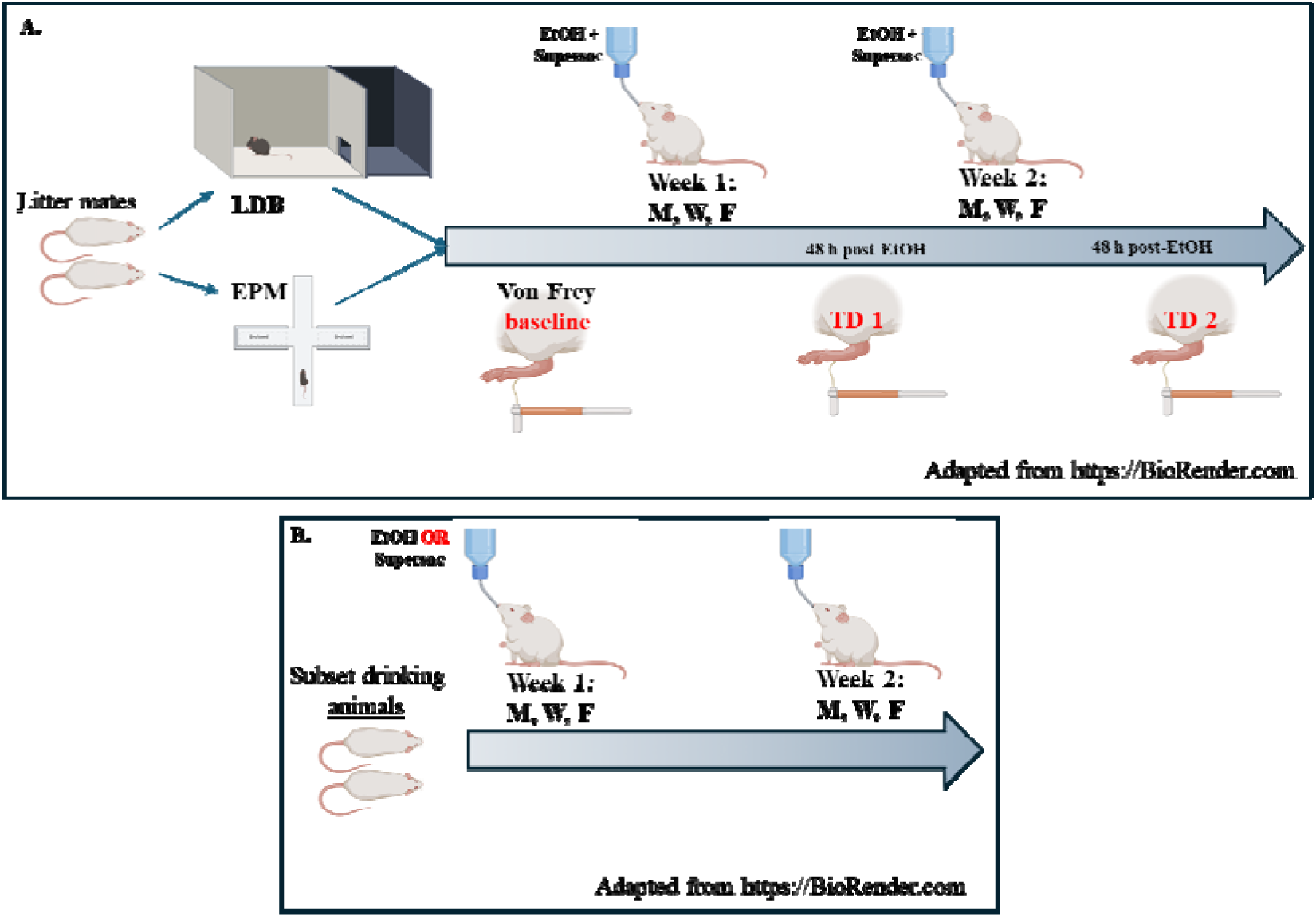
Experimental Timeline. **(A)** Outlines the original timeline, where different subsets of animals for each age group went through experimental testing. **(B)** Outlines the secondary timeline for additional drinking experiments, where different subsets of animals were tested on *EtOH only* and *super-sac only* experiments.

Animals were handled for two days before behavioral testing. On test day, cage mates were split and tested once in either the LDB or the EPM. LDB and EPM testing were video recorded and scored by a trained researcher blind to exposure conditions. All animals were then tested in a 30-minute limited access drinking paradigm with EtOH + SS solution. A baseline measurement of mechanical allodynia was taken directly after LDB/EPM behavioral testing and was repeated 48 hours after the 1^st^ week of drinking sessions (3^rd^ drinking session; Test Day 1 = TD1) and 48 hours after the 2^nd^ week of drinking sessions (6^th^ drinking session; Test Day 2 = TD2) (**Figure 1**). Two different subsets of animals were used for the second EtOH experiment and SS intake experiments; these animals did not go through LDB, EPM, or Von Frey testing.

### Statistical Analysis

All statistical analyses were performed using GraphPad 9 software (Prism). For behavior analysis, our *a priori* hypothesis was that there would be age- and sex- differences, based on previous reports of PAE differentially affecting EtOH intake and anxiety-like behaviors in males and females (Diaz et al., 2020; Rouzer et al., 2017). However, since this was a systematic ontogenetic assessment of these behaviors, all data collected from LDB and EPM were analyzed using a 2 (exposure: EtOH, air) x 2 (sex: male, female) analysis of variance (ANOVA) to determine if there in fact were sex differences at the various ages. Total EtOH intake was analyzed using a 2 (exposure: EtOH, air) x 2 (sex: male, female) ANOVA. In our *first* (EtOH + SS) and *second* (EtOH only) EtOH experiments, a main effect of sex or a sex x exposure interaction was observed in the total EtOH intake of most age groups (excluding juveniles in EtOH+SS and EtOH only experiments), therefore all drinking experiments were separated by sex and analyzed using either an unpaired t-test for total EtOH intake and BECs (effect of exposure) or a two-way repeated measures ANOVA: 2 (exposure) x 6 (session). A simple linear regression was computed for detecting the relationship between session 6 EtOH *+* SS intake and BECs. Von Frey data were analyzed using a two-way repeated measures 2 (exposure) x 2 (sex) ANOVA, which revealed sex as a main effect and/or interactions between sex and other variables; as a result, Von Frey data was separated by sex and analyzed using a two-way ANOVA: 2 (exposure) x 3 (Test Day). Sidak’s multiple comparison tests were used to determine specific group differences in the event of significance. A Grubb’s test was performed on LDB, EPM, and Von Frey data to detect a single outlier; however, this was not performed on data collected from drinking experiments to account for individual differences. Significance was defined as p ≤ 0.05. All data is presented as mean + the standard error of the mean (SEM).

## Results

### Light-Dark Box

In juvenile subjects (air males, n=7; air females, n=7; PAE males, n=8; PAE females, n=8), all behavioral measures in the LDB were unaffected by PAE (all p’s > 0.05; see **Table 1** for ANOVA results). Time spent in the light chamber (**Figure 2A**), egress latency (**Figure 2D**), and head pokes **(Figure 2G)** did not differ as a function of exposure or sex, with no interaction evident between these factors (all p’s > 0.05).

**Figure 2:**
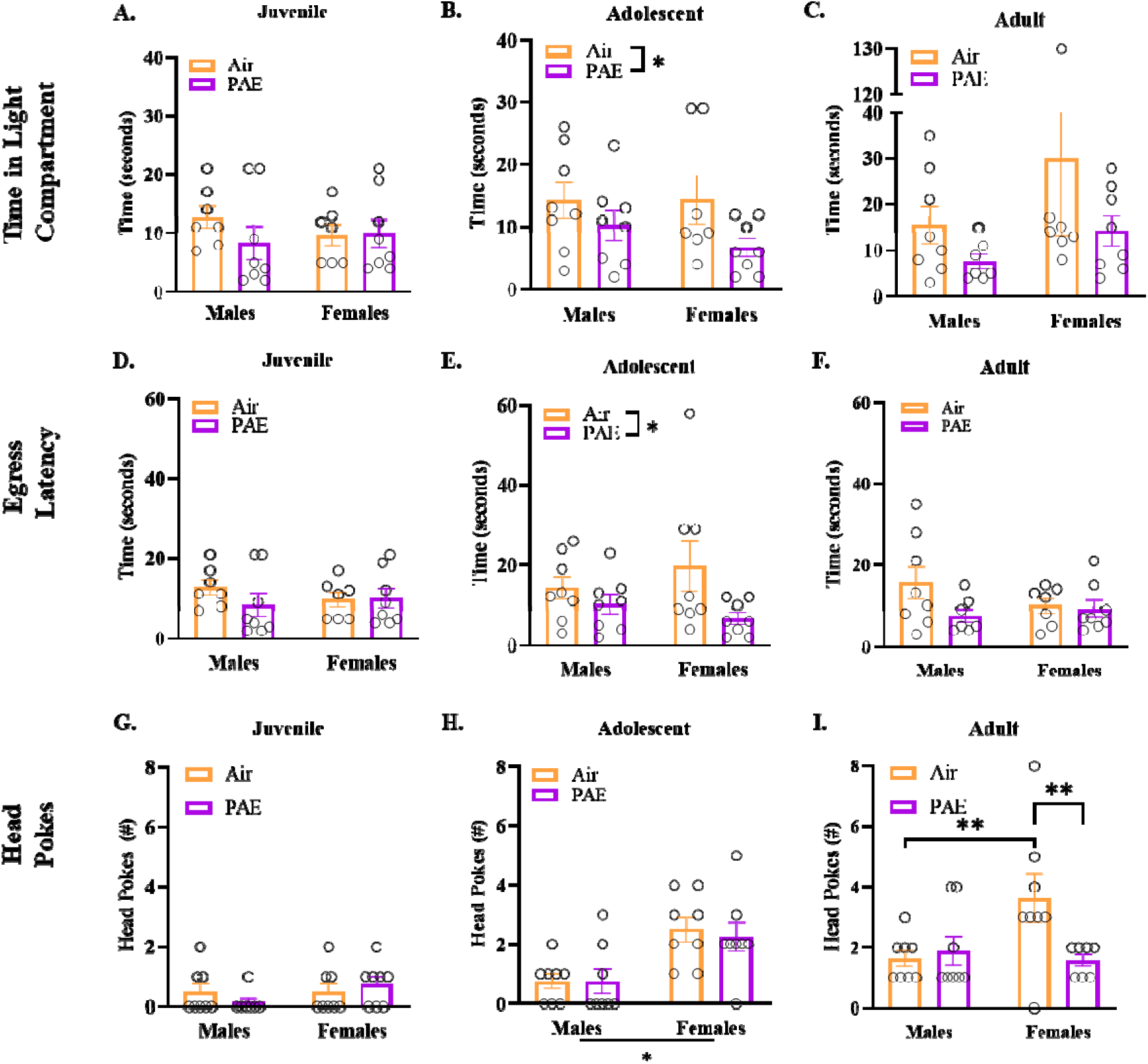
Light Dark Box (LDB). No differences were observed in male or female juveniles (n=7-8 per group) in **(A)** time spent in light compartment, **(D)** egress latency (time to enter dark compartment), or head pokes **(G)**. For adolescents (n=7-8 per group), there is a main effect of exposure observed in time spent in the light compartment **(B)**, with PAE animals spending less time in the light compartment. **(E)** PAE animals also have a shorter latency to enter the dark compartment. **(H)** No difference in adolescent head pokes. No differences were observed in adults (n=7-8 per group) in **(C)** times spent in light compartment, **(F)** egress latency. **(I)** sex x exposure interaction revealed adult air females performed more head pokes compared to air males and PAE counterpart. Two-way ANOVA was used to test for differences with a Sidak’s multiple comparison test when necessary. * denotes p ≤ 0.05. Bars represent standard error of the mean. PAE = Prenatal Alcohol Exposure.

**Table 1.**
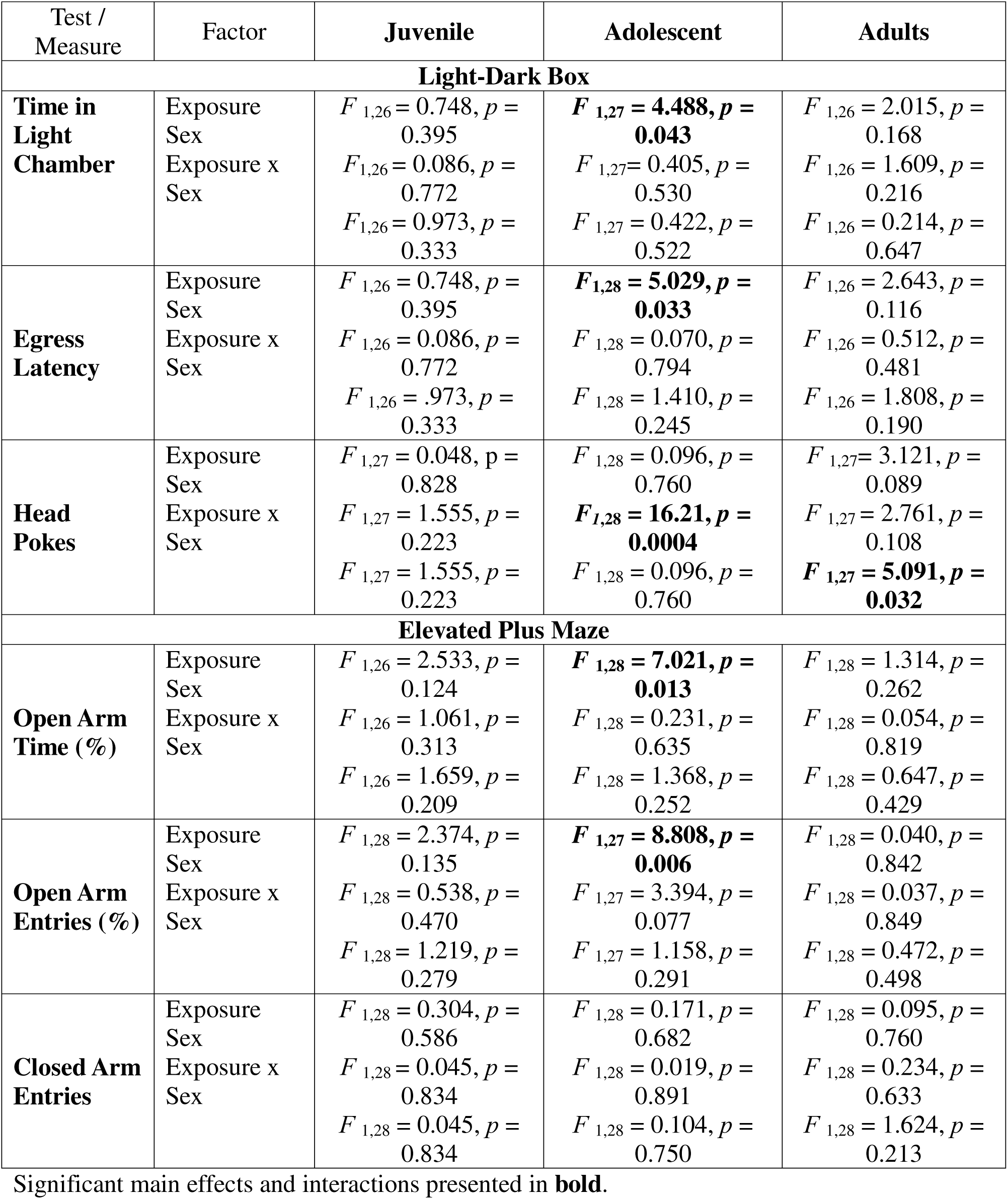
ANOVA results for anxiety-like behavioral measures presented in Figures 2 and 3.

In contrast, adolescent offspring (air males, n=8; air females, n=7; PAE males, n=8; PAE females, n=8) were affected by PAE (**Table 1**). PAE adolescents, regardless of sex, spent significantly less time in the light compartment than controls [F(1,27) = 4.488, p = 0.043], suggesting heightened anxiety-like behaviors (**Figure 2B**). Egress latency (**Figure 2E**) also significantly differed as a function of exposure [F(1,28) = 5.029, p = 0.033], with PAE animals demonstrating shorter latency to enter the dark compartment than air controls, likely driven by PAE females; however, there was no sex effect or exposure by sex interaction (p’s > 0.05). Head pokes significantly differed only as a function of sex [F(1,28) = 16.21, p = 0.0004], with females demonstrating more head pokes than males (**Figure 2H)**.

In adults (air males, n=8; air females, n=7; PAE males, n=7; PAE females, n=8), time spent in the light compartment (**Figure 2C**) and egress latency (**Figure 2F)** did not differ as a function of exposure or sex (p’s > 0.05). However, there was a significant interaction of exposure and sex for head pokes [F(1,27) = 5.091, p = 0.032; **Figure 2I**], with air females showing more head pokes than air males (*p* = 0.009) and PAE females (*p* = 0.009), suggesting that this behavioral measure was affected by PAE in female subjects, but not their male counterparts.

### Elevated Plus maze

To assess anxiety-like behavior across development in a slightly different context, a different set of animals were tested in the EPM, a standard assay of anxiety-like behavior in the field (Kraeuter et al., 2019).

Behavior of juveniles in the EPM was not affected by PAE (air males, n=8; air females, n=7-8; PAE males, n=7-8; PAE females, n=8). Percent open arm time (**Figure 3A**), percent open arm entries (**Figure 3D**), and number of closed arm entries (**Figure 3G**) did not differ as a function of exposure or sex (all p’s > 0.05; see **Table 1**). Risk assessment behavioral measures in the EPM (protected and unprotected head dips, protected and unprotected stretch attends) were also unaffected by PAE or sex in juvenile subjects (all p’s > 0.05; **Supplementary Table 1).**

**Figure 3:**
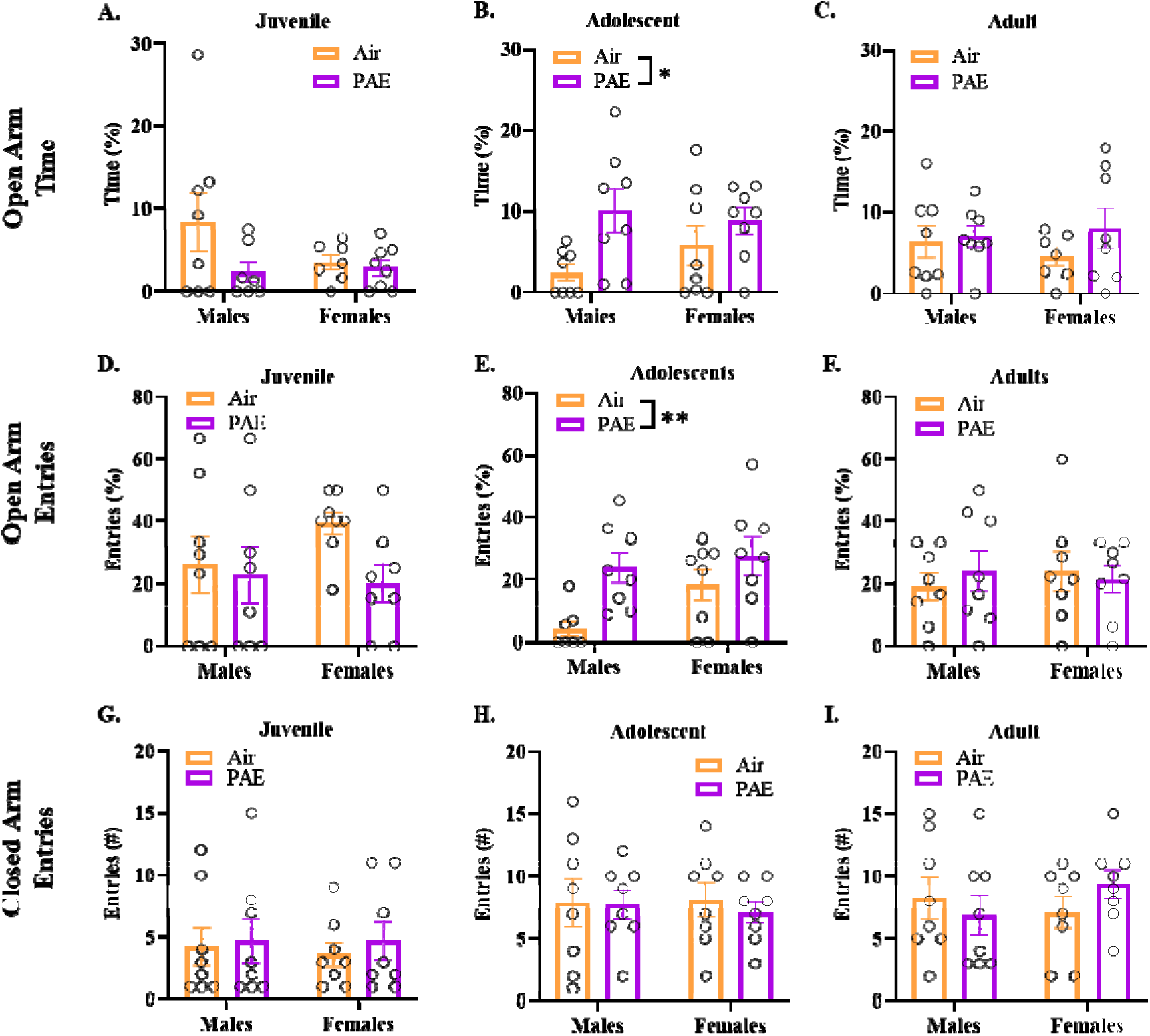
Elevated Plus Maze. No differences observed in juveniles (n=7-8 per group) in **(A)** percent time spent in the open arms, **(D)** percent of open arm entries or **(G)** closed arm entries. **(B)** Adolescent PAE animals spent more time in the open arms compared to controls (n=8 per group). **(E)** A main effect of exposure was observed in percent of open arm entries in adolescent animals. **(H)** No differences were observed in adolescent animals for closed arm entries. No differences were observed in adults (n=8 per group) in **(C)** percent time spent in open arms, **(F)** percent of open arm entries, or **(I)** number of closed arm entries. Two-way ANOVA was used to test for differences with a Sidak’s multiple comparison test when necessary. * denoted p ≤ 0.05. Bars represent standard error of the mean. PAE = Prenatal Alcohol Exposure.

In adolescents, anxiety-like behavior in the EPM was affected by PAE (**see Table 1**), with PAE adolescents, surprisingly, showing less anxiety-like responses than air controls (air males, n=8; air females, n=8; PAE males, n=8; PAE females, n=8). Specifically, PAE animals spent more time in the open arms [F(1,28) = 7.021, p = 0.013; **Figure 3B**] and demonstrated more open arm entries [F(1,27) = 8.808, p = 0.006; **Figure 3E**] than air controls, with these effects likely driven by PAE males. In contrast, closed arm entries were not affected by PAE in male or female adolescents (p > 0.05; **Figure 3H)**. Although protected head dips and stretch attends were unaffected by PAE in adolescent animals, PAE animals conducting a higher number of unprotected head dips [F(1,28) = 4.129, p = 0.052]. Unprotected stretch attends differed as a function of sex [F(1,28) = 6.725, p = 0.015], with adolescent females showing significantly more unprotected stretch attends than adolescent males (**Supplementary Table 1**).

In adults (air males, n=8; air females, n=7-8; PAE males, n=8; PAE females, n=8), percent open arm time (**Figure 3C**), percent open arm entries (**Figure 3F**), as well as closed arm entries (**Figure 3I**) were not affected by PAE and did not differ as a function of sex (all p’s > 0.05; **Table 1**). All measures of risk assessment behavior were unaffected by PAE in adult male and female subjects, with no sex differences evident at this age as well (all p’s > 0.05; **Supplementary Table 1).**

### EtOH + SS

To assess PAE’s effect on subsequent EtOH intake across ontogeny, one day after LDB/EPM testing and collection of baseline Von Frey measures (see below for results on Von Frey), animals were tested in a 30-minute limited access drinking paradigm for six sessions over two weeks with access to 10% EtOH + SS solution.

In juveniles (n=15 per group), we observed no significant sex, exposure, or exposure by sex differences in total EtOH intake (all p’s > 0.05; **Figure 4A**). However, assessment of EtOH + SS intake across sessions in juvenile males revealed an exposure by session interaction [F(5,150) = 3.224, p = 0.0327; see **Table 2**], with PAE males consuming less EtOH + SS during session six than air controls (p = 0.027; **Figure 4B**). In juvenile females, intake of EtOH + SS was not affected by PAE and did not change across sessions (p’s > 0.05; **Figure 4C**). BECs assessed after the last drinking session did not differ as a function of exposure in either sex (p’s > 0.05; **Table 2**, **Figure 4D**), but were significantly correlated with EtOH intake in air males, air females, and PAE females, with no significant correlation evident in PAE males (**Supplementary Table 2**).

**Figure 4:**
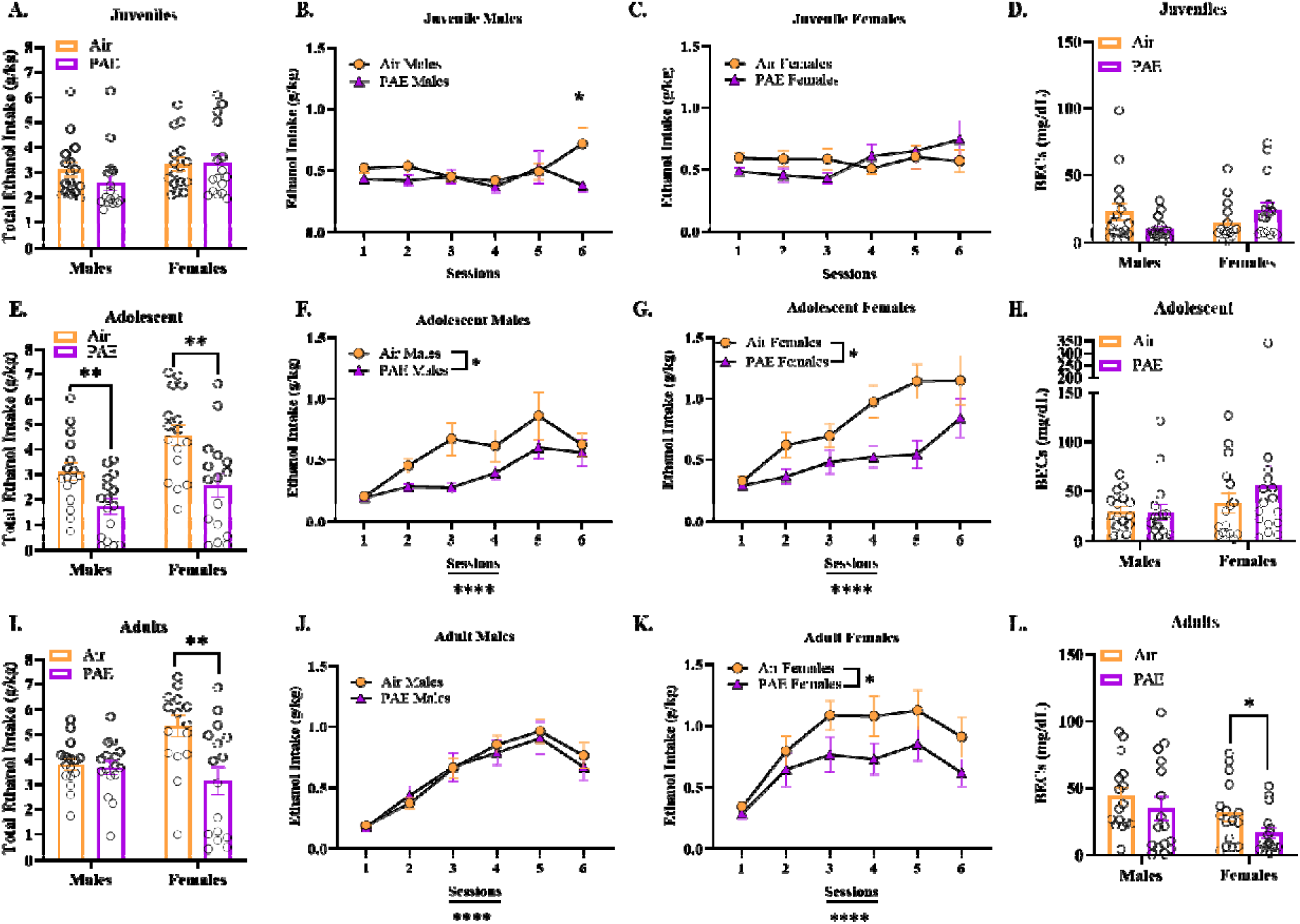
*Ethanol + Super-sac* Intake. A main effect of sex was found in total EtOH intake in most groups, therefore data was split by sex and analyzed separately. **(A)** No differences observed in juvenile total intake **(B)** Exposure by session interaction, where juvenile air males consumed more EtOH on session 6. **(C)** No effect of exposure or session observed in juvenile females **(D)**. No differences observed in session 6 BECs for juvenile male or females (n=16 per group). **(E)** Adolescent PAE consumed significantly less total EtOH compared to air counterparts (n=16 per group). PAE males **(F)** and females **(G)** consumed less EtOH compared to air animals across all sessions. **(H)** No differences were observed in session 6 BECs for adolescent males or females (n=16 per group). **(I)** No differences were observed in adult male total intake, but PAE females consumed less EtOH overall (n=16 per group). **(J)** No differences observed in males. **(K)** PAE females consumed less EtOH compared to air controls (n=16 per group). **(L)** No differences observed in BECs for adult males. PAE females had lower BECs compared to air control females (n=16 per group). T-test was used to determine differences in EtOH total intake and BECs. Repeated measures ANOVA was used to test for differences across sessions with a Sidak’s multiple comparison test when necessary. * denoted p ≤ 0.05. Bars represent standard error of the mean. BECs = Blood Ethanol Concentrations. EtOH = ethanol. PAE = Prenatal alcohol exposure.

**Table 2.**
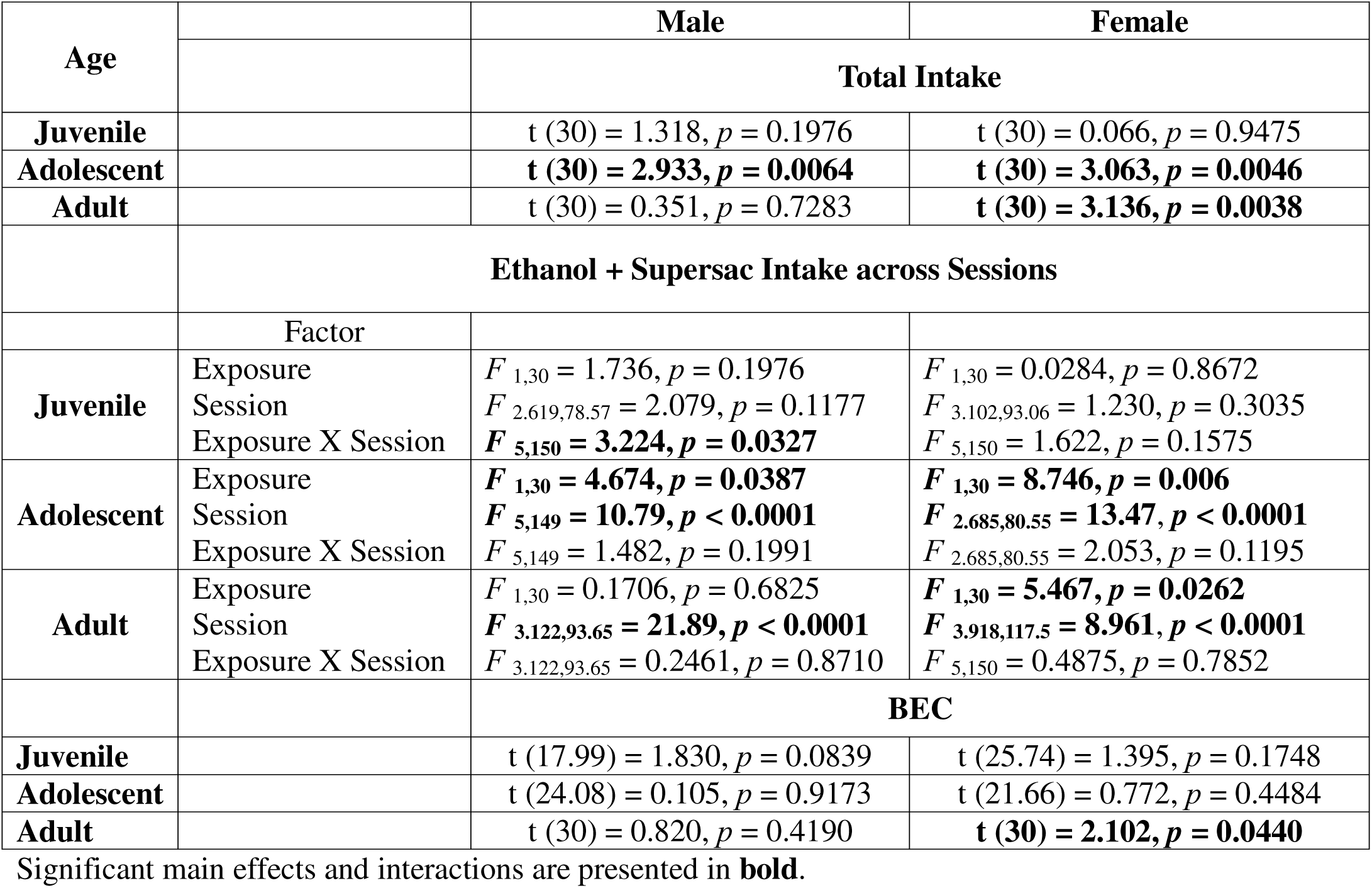
ANOVA results for ethanol + supersac intake and blood ethanol concentration (BEC) presented in Figure 4.

In adolescents (n=16 per group), total EtOH + SS intake differed as a function of exposure [F(1,60) = 17.62, p < 0.0001] and sex [F(1,60) = 8.264, p = 0.0056], with PAE adolescents drinking significantly less than their air counterparts and females drinking more than males (**Table 2**, **Figure 4E)**. An effect of session was observed in male [F(5,149) = 10.79, p < 0.000] and female [F(2.685,80.55) = 13.47, p < 0.0001] adolescents, with adolescent animals increasing intake over sessions (p < 0.05), yet PAE males [F(1,30) = 4.674, p = 0.0387] and females [F(1,30) = 8.746, p = 0.006] consumed significantly less than their control counterparts (see **Table 2**, **Figure 4F, 4G**), with no session by exposure interactions (p > 0.05). BECs, however, were not affected by exposure (p’s > 0.05; **Table 2**, **Figure 4H**). BECs assessed after session 6 were significantly correlated with EtOH + SS intake in PAE males and air females, with no significant correlations evident in air males and PAE females (**Supplementary Table 2**).

In adult males and females (n=16 per group), although there was no main effect of sex on total EtOH intake (p > 0.05), there was a significant main effect of exposure [F(1,60) = 8.663, p = 0.0046] and a exposure by sex interaction [F1(1,60) = 6.862, p = 0.0111]. While we did not observe any differences in the total EtOH consumption for males (p > 0.05), air females consumed more EtOH overall when compared to PAE counterparts (p = 0.0038; **Figure 4I**, **Table 2**). EtOH + SS consumption increased across drinking sessions in males [F(3.122,93.65) = 21.89, p < 0.0001] and females [F(3.918,117.5) = 8.961, p < 0.0001], with PAE females [F(1,30) = 5.467, p = 0.0262], but not PAE males (p > 0.05) ingesting less EtOH than their air counterparts (**Table 2**, **Figure 4J, 4K).** BECs did not differ as a function of exposure in males (p > 0.05; **Figure 4L**), however, PAE females displayed significantly lower BECs compared to their counterparts [t(30) = 2.102, p = 0.0440]. Session 6 EtOH + SS intake was positively correlated with BECs in air males, PAE males, air females, and PAE females (**Supplementary Table 2**).

### Unsweetened Ethanol Intake

Pre-clinical and clinical research have shown that individuals with a history of PAE tend to consume more alcohol than those not exposed (Baer et al., 1998; Baer et al., 2003; Diaz et al., 2020; Gaztanaga et al., 2020), yet we observed a reduction in intake in PAE animals when using sweetened EtOH (EtOH + SS). Therefore, to assess if moderate G12 PAE affects unsweetened EtOH intake differently, using a separate cohort of animals we tested intake of unsweetened 10% EtOH using the same 30-minute limited access drinking paradigm.

In juveniles, total EtOH intake in PAE animals was significantly higher than EtOH intake demonstrated by air controls [F(1,30] = 19.85, p = 0.0001], with no sex effects (p > 0.05) or sex by exposure interaction (p > 0.05) evident at this age **(Table 3**, **Figure 5A).** In juvenile males (air: n=10; PAE: n=8), EtOH intake differed as a function of exposure [F(1,16) = 8.018, p = 0.0120] and session [F(5,80) = 16.74, p < 0.0001], with a significant exposure by session interaction evident as well [F(5,80) = 2.774, p = 0.0231; see **Table 3**, **Figure 5B**]. Post-hoc analysis showed that during session one PAE males consumed more EtOH than air controls (*p* = 0.0005), with EtOH intake gradually decreasing across sessions. In juvenile females (n=8 per prenatal exposure condition), there was an effect of exposure [F(1,14) = 14.11, p = 0.0021] and session [F(4.049,56.69) = 5.134, p = 0.0013; **Table 3**, **Figure 5C**], with females decreasing intake across sessions.

**Figure 5:**
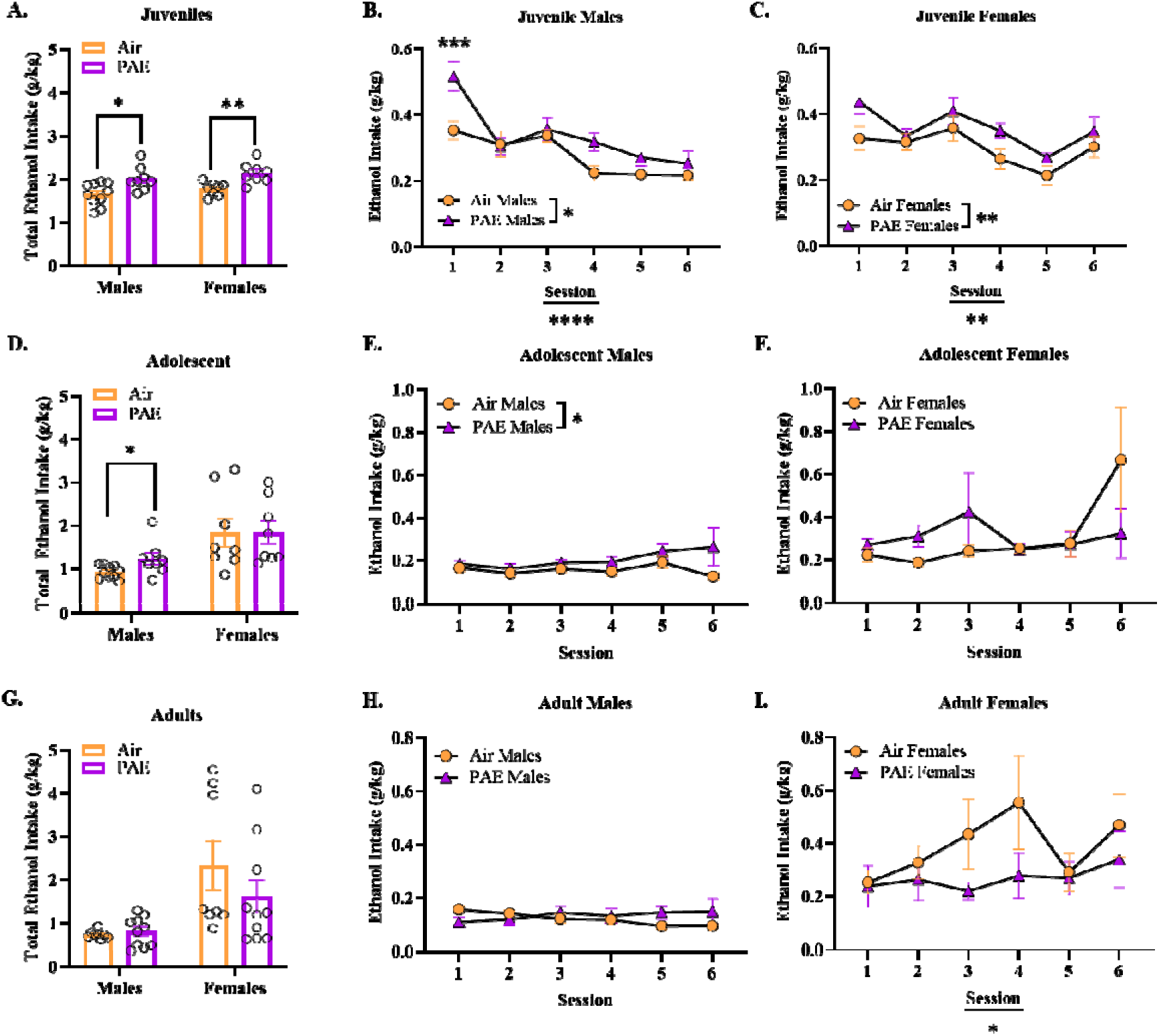
*Ethanol Only* Intake. Data was split by sex and analyzed separately. **(A)** Air animals consumed less overall EtOH compared to PAE animals (n= 8-10 per group). **(B)** There was a main effect of exposure across male drinking sessions, and an exposure x session interaction with PAE males consuming more on session 1. **(C)** PAE females consumed more than air counterparts across sessions. **(D)** Adolescent PAE males consumed more overall EtOH when compared to air counterparts, with no differences in female total intake (n=8-10 per group). **(E)** PAE males consumed more EtOH compared to air males across sessions, **(F)** with no differences observed in females across sessions. No differences were observed in **(G)** adult total EtOH intake (n= 8-10 per group), or across sessions in **(H)** males or **(I)** females. T-test was used to determine differences in EtOH total intake. Repeated measures ANOVA was used to test for differences across sessions with a Sidak’s multiple comparison test when necessary. * denoted p ≤ 0.05. Bars represent standard error of the mean. EtOH = ethanol. PAE = Prenatal alcohol exposure.

**Table 3.**
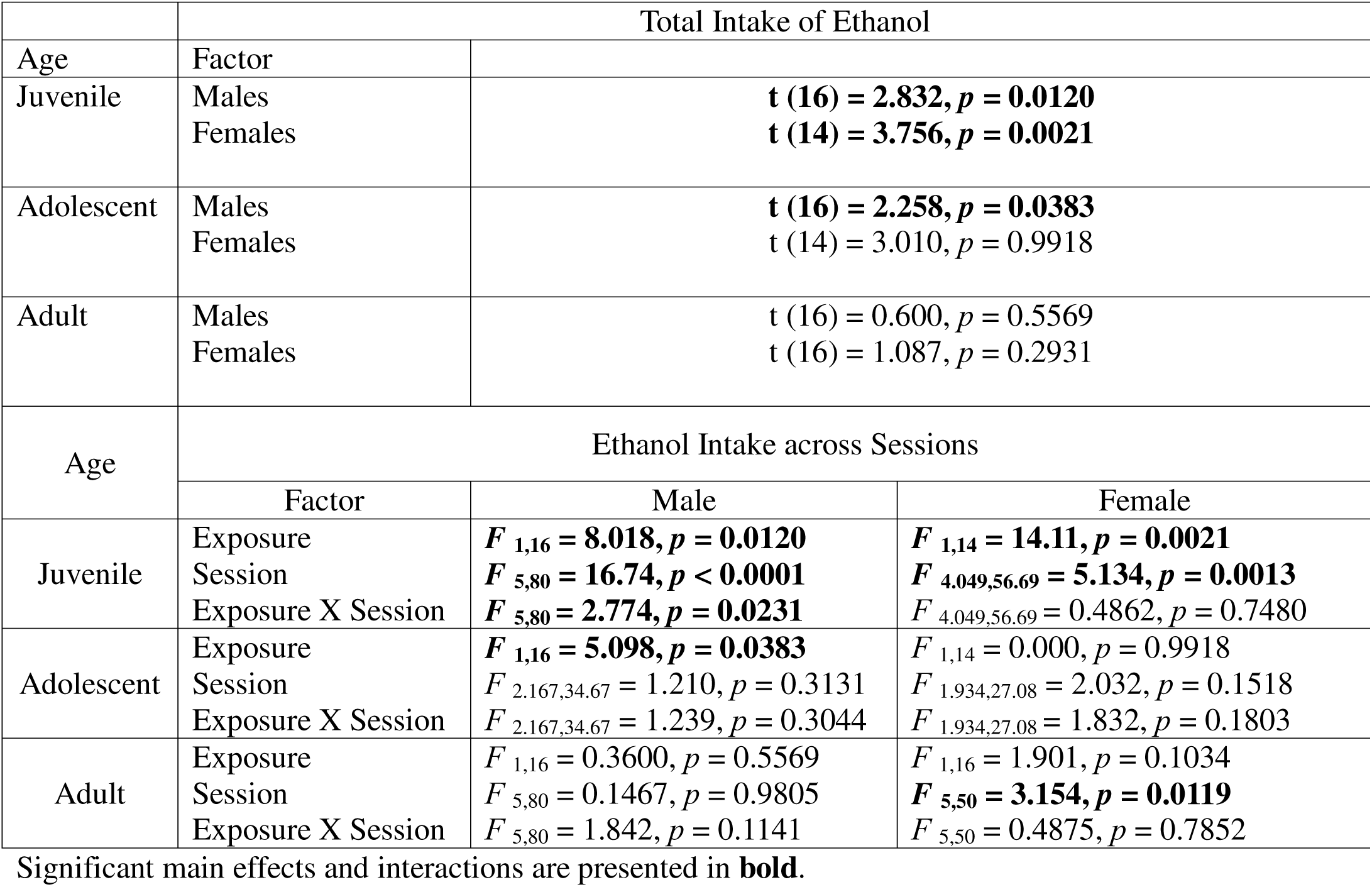
ANOVA results for unsweetened ethanol intake presented in Figure 5.

|  | Total Intake of Ethanol |  |  |
| --- | --- | --- | --- |
| Age | Factor |  |  |
| Juvenile | Males<br>Females | **t (16) = 2.832, p = 0.0120**<br>**t (14) = 3.756, p = 0.0021** |  |
| Adolescent | Males<br>Females | **t (16) = 2.258, p = 0.0383**<br>t (14) = 3.010, p = 0.9918 |  |
| Adult | Males<br>Females | t (16) = 0.600, p = 0.5569<br>t (16) = 1.087, p = 0.2931 |  |
| Age | Ethanol Intake across Sessions |  |  |
|  | Factor | Male | Female |
| Juvenile | Exposure<br>Session<br>Exposure X Session | **F<sub>1,16</sub> = 8.018, p = 0.0120**<br>**F<sub>5,80</sub> = 16.74, p < 0.0001**<br>**F<sub>5,80</sub> = 2.774, p = 0.0231** | **F<sub>1,14</sub> = 14.11, p = 0.0021**<br>**F<sub>4.049,56.69</sub> = 5.134, p = 0.0013**<br>F<sub>4.049,56.69</sub> = 0.4862, p = 0.7480 |
| Adolescent | Exposure<br>Session<br>Exposure X Session | **F<sub>1,16</sub> = 5.098, p = 0.0383**<br>F<sub>2.167,34.67</sub> = 1.210, p = 0.3131<br>F<sub>2.167,34.67</sub> = 1.239, p = 0.3044 | F<sub>1,14</sub> = 0.000, p = 0.9918<br>F<sub>1.934,27.08</sub> = 2.032, p = 0.1518<br>F<sub>1.934,27.08</sub> = 1.832, p = 0.1803 |
| Adult | Exposure<br>Session<br>Exposure X Session | F<sub>1,16</sub> = 0.3600, p = 0.5569<br>F<sub>5,80</sub> = 0.1467, p = 0.9805<br>F<sub>5,80</sub> = 1.842, p = 0.1141 | F<sub>1,16</sub> = 1.901, p = 0.1034<br>**F<sub>5,50</sub> = 3.154, p = 0.0119**<br>F<sub>5,50</sub> = 0.4875, p = 0.7852 |

Significant main effects and interactions are presented in **bold**.

In adolescents, females consumed significantly more total EtOH compared to males [F(1,30) = 13.52, p = 0.0009]. Furthermore, within males, PAE males demonstrated significantly higher total EtOH intake relative to air males (p = 0.0383), with no differences in females (p > 0.05; **Table 3**, **Figure 5D**). These effects were also apparent across sessions, with a significant increase in EtOH intake in adolescent PAE males relative to air males (air: n=10; PAE: n=8; F(1,16) = 5.098, p = 0.0383), with no main effects of session or exposure by session interactions (p’s > 0.05; **Table 3**, **Figure 5E**). Conversely, intake of EtOH in adolescent females (n=8 per group) was not affected by exposure and did not change across sessions (p’s > 0.05; **Table 3**, **Figure 5F**).

In adults there was a main effect of sex on total EtOH intake, with females consuming more than males [F(1,32) = 12.88, p = 0.0011], however, there were no effects of exposure on total EtOH intake in either sex (p’s > 0.05; **Table 3**, **Figure 5G**). When examining EtOH intake across sessions, intake in adult males (air: n=8; PAE: n=10) was not affected by either exposure or session (p’s > 0.05; **Table 3**, **Figure 5H**). Conversely, there was an effect of session in adult females [air: n=8; PAE: n=10; F(5,50) = 3.154, p = 0.0119], with EtOH intake increasing across sessions (**Table 3**, **Figure 5I**).

### Supersac (SS) Intake

To further parse apart the effects we observed in EtOH drinking experiments, a separate cohort of animals was tested in the 30-minute limited access drinking paradigm with access to SS solution only. SS intake was not affected by PAE in males or females at any age (all p’s > 0.05 (see **Table 4**).

**Table 4.** Intake of supersac in g/kg across sessions (mean± SEM) and ANOVA results.

| Age | Session / Statistics | Air Male | PAE Male | Air Female | PAE Female |
| --- | --- | --- | --- | --- | --- |
| Juvenile |  | n= 8 | n = 10 | n= 8 | n = 10 |
|  | 1 | 2.976 ± 1.419 | 1.018 ± 0.406 | 3.056 ± 1.034 | 1.958 ± 0.928 |
|  | 2 | 3.647 ± 2.916 | 2.193 ± 0.921 | 2.353 ± 0.989 | 6.270 ± 2.431 |
|  | 3 | 2.909 ± 0.929 | 3.035 ± 1.811 | 4.243 ± 1.621 | 4.543 ± 2.561 |
|  | 4 | 3.490 ± 2.340 | 2.225 ± 1.310 | 4.590 ± 2.115 | 9.254 ± 4.079 |
|  | 5 | 2.377 ± 2.173 | 1.936 ± 1.392 | 1.942 ± 1.582 | 4.178 ± 2.231 |
|  | 6 | 2.232 ± 1.269 | 0.621 ± 0.385 | 4.436 ± 2.474 | 7.684 ± 3.907 |
|  | Total Intake | t (12.25) = 0.9524, *p* = 0.3593 |  | t (16) = 0.9808, *p* = 0.3413 |  |
| Adolescent | Session / Statistics | Air Male | PAE Male | Air Female | PAE Female |
|  |  | n= 10 | n = 10 | n= 10 | n = 8 |
|  | 1 | 1.531 ± 0.618 | 3.107 ± 2.740 | 4.261 ± 1.154 | 2.460 ± 1.028 |
|  | 2 | 4.111 ± 1.676 | 12.06 ± 5.916 | 10.88 ± 4.254 | 5.308 ± 3.391 |
|  | 3 | 6.943 ± 3.447 | 12.79 ± 7.913 | 18.58 ± 5.941 | 7.981 ± 4.122 |
|  | 4 | 4.129 ± 2.561 | 11.27 ± 4.705 | 15.93 ± 8.836 | 2.971 ± 1.756 |
|  | 5 | 5.007 ± 3.304 | 6.323 ± 4.707 | 25.42 ± 9.614 | 3.876 ± 2.745 |
|  | 6 | 4.032 ± 2.028 | 9.067 ± 5.613 | 15.24 ± 8.527 | 11.03 ± 7.769 |
|  | Total Intake | t (13.23) = 0.8672, *p* = 0.4013 |  | t (16) = 1.596, *p* = 0.130 |  |
| Adult | Session / Statistics | Air Male | PAE Male | Air Female | PAE Female |
|  |  | n= 10 | n = 10 | n= 10 | n = 8 |
|  | 1 | 0.586 ± 0.386 | 2.469 ± 1.627 | 6.934 ± 3.771 | 5.459 ± 2.782 |
|  | 2 | 2.516 ± 1.883 | 1.035 ± 0.673 | 5.389 ± 5.368 | 0.015 ± 0.008 |
|  | 3 | 2.519 ± 1.683 | 0.013 ± 0.007 | 12.91 ± 5.375 | 7.127 ± 6.963 |
|  | 4 | 7.627 ± 4.056 | 0.885 ± 0.865 | 19.09 ± 9.066 | 12.16 ± 7.886 |
|  | 5 | 4.339 ± 4.292 | 0.483 ± 0.469 | 14.21 ± 9.266 | 0.577 ± 0.443 |
|  | 6 | 2.492 ± 2.399 | 0.395 ± 0.375 | 13.20 ± 8.895 | 2.451 ± 2.397 |
|  | Total Intake | t (18) = 1.496, *p* = 0.1520 |  | t (10.91) = 1.202, *p* = 0.2546 |  |

Significant main effects and interactions are presented in **bold**

### Von Frey Test

To evaluate the effect of PAE on mechanical sensitivity, mechanical thresholds were determined using Von Frey filaments, reported as target force (g). Baseline assessments were conducted immediately after anxiety behavior testing and prior to assessments of EtOH + SS intake. Two additional testing days occurred 48 hours after the 3^rd^ and 6^th^ EtOH + SS session to examine the effect of postnatal EtOH consumption on mechanical thresholds as a function of PAE. A two-way repeated measures ANOVA revealed an effect of sex in juveniles [F(1,59) = 9.450, *p* = 0.003], adolescents [F(1,60) = 10.09, *p* = 0.002], and adults [F(1,60) = 13.27, *p* = 0.006]. Therefore, data was split by sex, analyzed and presented separately for males and females across ontogeny (see **Table 5** for ANOVA results).

**Table 5.**
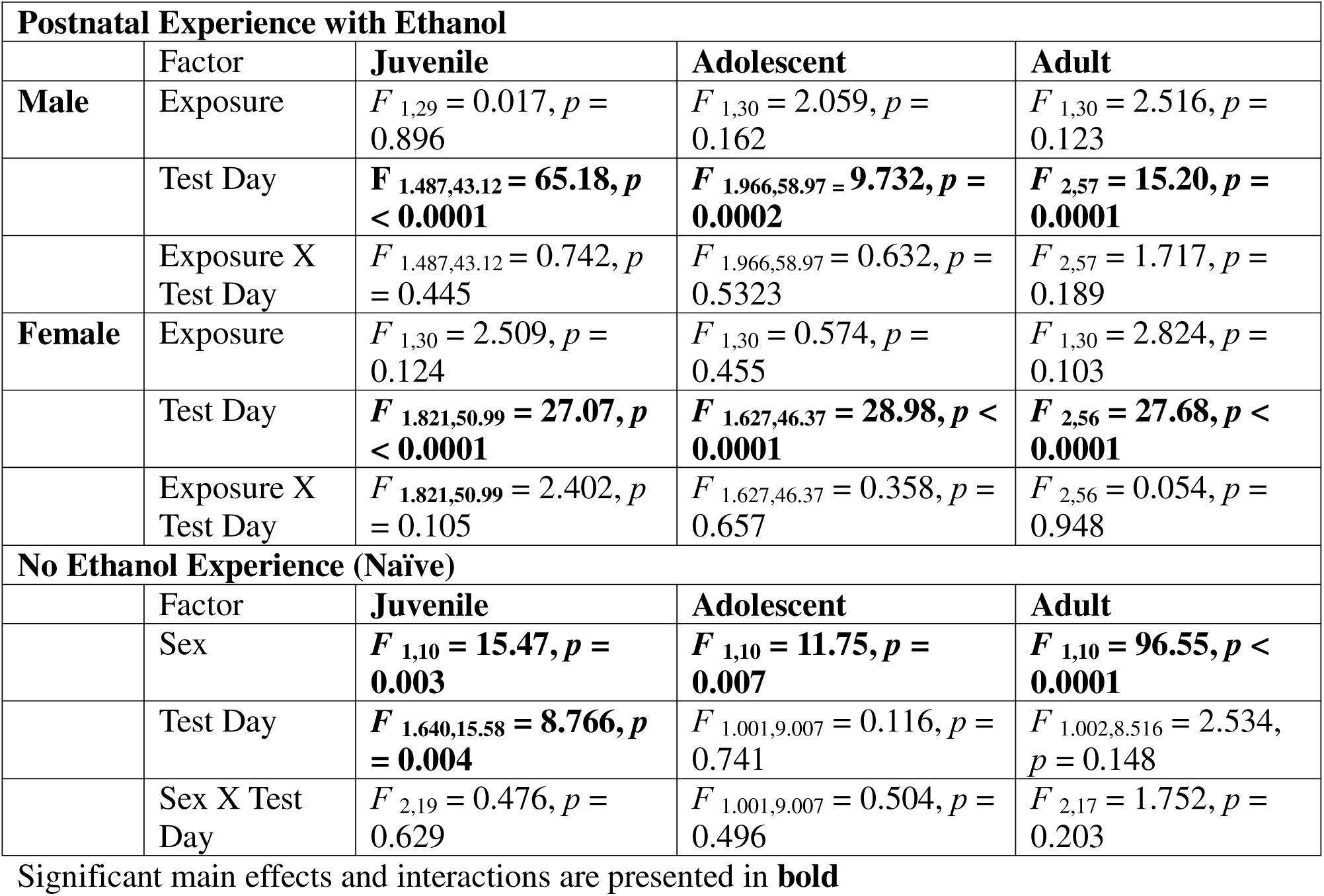
ANOVA results for mechanical thresholds determined with Von Frey filaments and presented on. **Figure 6**.

In juvenile males (air: n=15; PAE: n=16), mechanical thresholds did not differ as a function of exposure (p > 0.05), but there was an effect of test day (**Figure 6A**), in that thresholds decreased on both test days relative to baseline [F(1.487,43.12) = 65.18, p < 0.0001]. In adolescent males (n=16 per group), mechanical thresholds also differed only as a function of test day (**Figure 6C**), in that thresholds declined relative to baseline [F(1.966,58.97) = 9.732, p = 0.0002], although there was no effect of PAE (p > 0.05). In adult males (n =16 per group), there was also no effect of exposure on mechanical thresholds (p > 0.05), however, similar to juveniles and adolescents, there was an effect of test day [F(2,57) = 15.20, p = 0.0001], where thresholds decreased on test day 2 and 3 relative to baseline (**Figure 6E**). Together, these findings suggest that mechanical sensitivity increased in male subjects following EtOH intake regardless of prenatal exposure condition.

**Figure 6:**
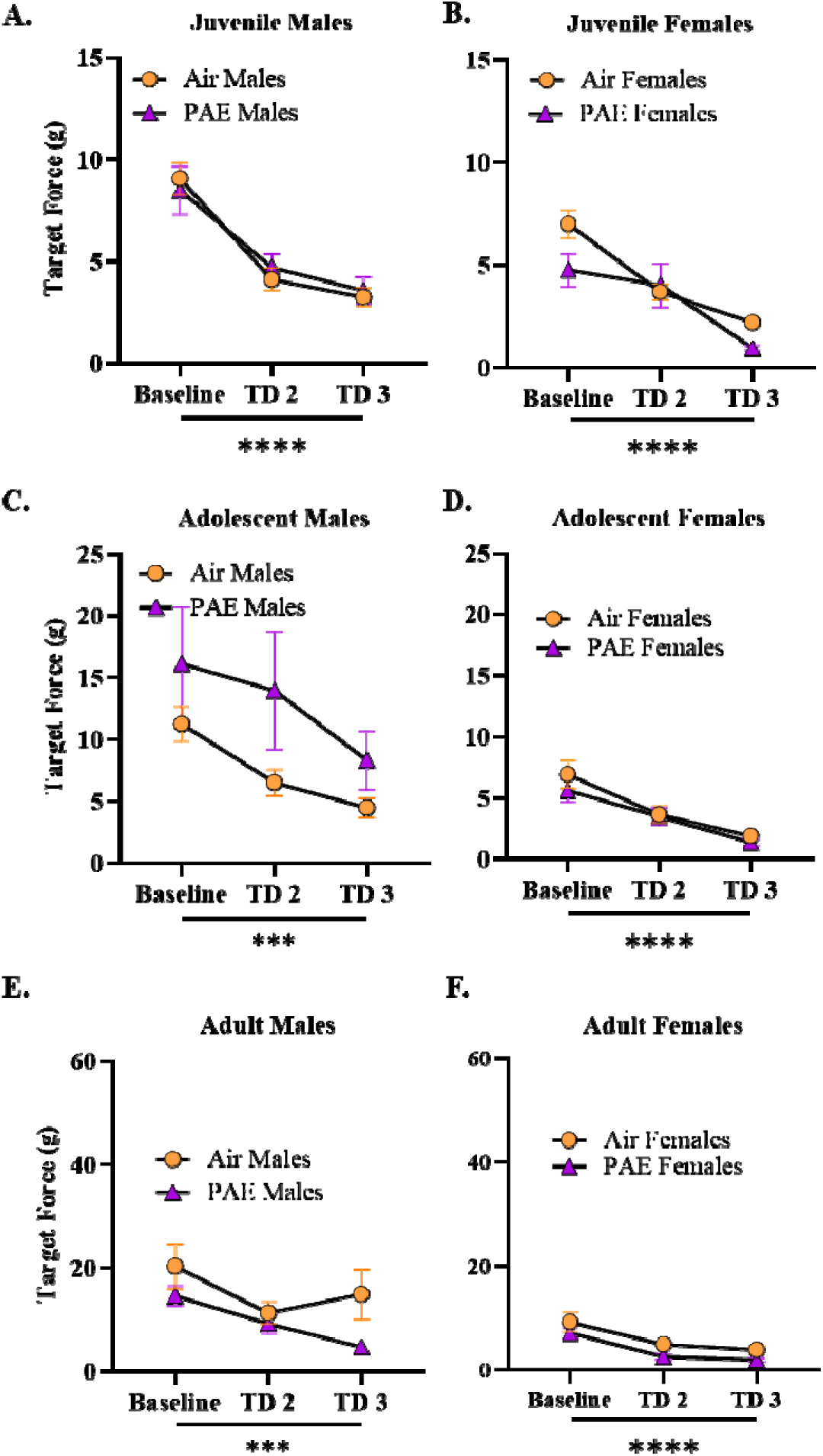
Von Frey. A two-way ANOVA revealed an effect of sex, therefore all data is split by sex. **(A-F)** There was no effect of exposure on mechanical thresholds in juveniles (n=15-16 per group), adolescents (n=16 per group), or adults (n=16 per group). **(A-F)** In each sex for every age group there was an effect of test day, where mechanical thresholds decreased over time. Repeated measures ANOVA was used to test for differences across test days with a Sidak’s multiple comparison test when necessary. * denoted p ≤ 0.05. Bars represent standard error of the mean. PAE = Prenatal alcohol exposure.

Similarly, in juvenile females (n=16 per group), there was only an effect of test day (**Figure 6B**), with thresholds decreasing across testing days [F(1.821,50.99) = 27.07, p < 0.0001]. However, there was no effect of exposure and no exposure by test day interaction found in juvenile female mechanical thresholds (p’s > 0.05; see **Table 5**). In adolescent females (n=16 per group), thresholds also differed as a function of test days (**Figure 6D**), with thresholds on test day 2 and 3 decreasing relative to baseline [F(1.627,46.37) = 28.98, p < 0.0001] with no effect of exposure or exposure by test day interaction (p’s > 0.05). In adult females (n=16 per group), mechanical thresholds differed as a function of test day [F(2,56) = 27.68, p < 0.0001; **Figure 6F**), but not of exposure (p > 0.05; **Table 5**). Together, these findings suggest that females also developed an increase in mechanical sensitivity as a function of EtOH intake, regardless of prenatal exposure condition and age.

Given the observed decreases in mechanical thresholds relative to baseline, indicative of enhanced mechanical sensitivity, it was important to determine whether the development of mechanical sensitivity evident in all groups was related to the EtOH consumption the animals were experiencing between the von Frey testing, and not an effect of repeated testing. To address this issue, we examined mechanical thresholds in EtOH-naïve animals (no pre- or postnatal EtOH exposures) using the same experimental timeline. In naïve juvenile animals (n=6 per group), we observed an effect of sex [F(1,10) = 15.47, p = 0.003] and test day [F(1.640,15.58) = 8.766, p = 0.004) on mechanical thresholds, but no interaction (p < 0.05; **Table 5**, **Supplementary Figure 1A**). Interestingly, thresholds increased in EtOH-naïve juvenile animals at test day 3 relative to baseline and test day 2, indicating less sensitivity over time. In EtOH-naïve adolescent animals, thresholds did not change across days (p > 0.05) and were significantly lower in females than males, as evidenced by a main effect of sex [F(1,10) = 11.75, p = 0.007; **Supplementary Figure 1B**]. In EtOH-naïve adults (n=6 per group), mechanical thresholds differed as a function of sex as well [F(1,10) = 96.55, p < 0.0001; **Table 5, Supplementary Figure 1C**], but not test day, suggesting that repeated testing did not affect thresholds.

## Discussion

A goal of this study was to investigate whether a single moderate PAE affects anxiety-like behaviors, EtOH intake, and mechanical sensitivity across ontogeny and to test whether PAE-associated changes are sex-dependent. Due to known dynamic and rapid alterations that occur during adolescence, we tested juvenile, adolescent, and adult males and females. Adolescence is characterized by substantial behavioral, hormonal, and neural changes (Spear, 2000, 2004, 2013), and in the current study, PAE adolescent animals, but not their juvenile or adult counterparts, exhibited increased anxiety-like behaviors in the LDB, with females likely more affected than males. In contrast, control adolescent males demonstrated heightened anxiety-like behavior in the EPM, with PAE decreasing this anxiety-like responding. We hypothesized that PAE would result in an increase in subsequent EtOH intake, yet, surprisingly, adolescent PAE males and females, as well as adult PAE females consumed substantially less sweetened EtOH compared to controls. In contrast, juvenile males and females, as well as adolescent PAE males consumed more unsweetened EtOH compared to air controls, with no differences observed in *SS only* intake. However, intake of unsweetened EtOH was extremely low, especially in adolescents and adults. Finally, although mechanical thresholds were not affected by PAE, thresholds progressively decreased after EtOH consumption across ontogeny, suggesting the emergence of mechanical sensitivity. Taken together, these data demonstrate that effects of PAE are more evident during adolescence, with sex-specific effects of PAE depending on age at testing.

This study revealed that PAE animals demonstrated heightened anxiety-like behavior in the LDB when tested during adolescence, an effect we have observed before (Rouzer et al., 2017). Although, in the present study, females tested at P45-55 were affected more than their male counterparts, whereas previously this PAE effect was seen in males tested at P40-42 (Rouzer et al., 2017). This suggests that age at testing may play a role in sex-dependent anxiety-like alterations associated with PAE, highlighting the dynamic effects of PAE across development. Interestingly, the main behavioral anxiety-related measure in the LDB test (i.e. time in light compartment) was not affected by PAE in juveniles or adults, suggesting that adolescence is a critical developmental period for the expression of anxiety-like behavior in the LDB in PAE offspring.

In general, PAE did not affect EPM behavior of juveniles or adults, while PAE adolescents showed less anxiety-like behavior in the EPM than air controls, with this effect of PAE driven predominantly by males. Increases in exploratory behaviors have been reported in late adolescent males following moderate PAE throughout the entire gestation period (Brolese et al., 2014) as well as in females with a history of PAE through liquid EtOH diet (Brolese et al., 2014; Osborn et al., 1998). While in the Brolese et al. study, controls spent about 20% of time in open arms, in the Osborn et al. study exploration of the open arms by controls was extremely low. Similarly, in the present study, adolescent control males demonstrated high levels of anxiety, with approximately half of the animals not showing open arm exploration, indexed via percent open arm time and entries. This finding was rather surprising given previous research in our lab that showed adolescents being less anxious in the EPM than adults (Przybysz et al., 2020). Previous research has shown that pretest manipulations affect exploration of the open arms in adult, but not adolescent rats, with a 30-min social isolation in a novel environment substantially increasing percent open arm time and entries (Doremus-Fitzwater et al., 2009). In the present study, animals were transferred to a novel cage and isolated for 1 hour prior to testing. Thus, it is possible that air control males, in contrast to PAE adolescent males, were not responsive to this pre-test manipulation. It is worth noting that since pretest manipulations can differentially affect open arm exploration in adolescent and adult rats, the EPM may not be the most suitable test for assessment of anxiety-like behavioral alterations associated with PAE across the lifespan, as previously observed by our lab (Przybysz et al., 2020). Alternatively, when utilizing the EPM, it should be used in parallel with other anxiety assays that are not as susceptible to pretest manipulations or age-dependent effects.

In the first EtOH intake experiment, we expected that PAE animals would consume more sweetened EtOH (EtOH + SS) than their control counterparts, as we previously found an increase in intake of EtOH sweetened with SS in adult offspring following a heavy dose G12 PAE that produced BECs > 300mg/dL (Diaz et al., 2020). Surprisingly, we found that PAE actually reduced EtOH + SS intake in adolescents of both sexes and adult females. Interestingly, when presented with an unsweetened EtOH solution, juvenile animals and adolescent males consumed more EtOH compared to their counterparts, however, intake of unsweetened EtOH was substantially lower than that of EtOH solution with SS, with decreases in intake becoming more evident across age. This finding suggests that the sweetener made EtOH more palatable for older animals than their less mature counterparts.

Multiple preclinical studies have reported PAE-associated increases in EtOH intake at varied ages ranging from P14 – P80 (Abate et al., 2008; Chotro and Arias, 2003; Fabio et al., 2015; Fabio et al., 2013; Gore-Langton and Spear, 2019); however, in all of these studies, the prenatal exposure occurred on G17-G20. The level and timing of exposure is a crucial component when examining maladaptive behaviors seen in individuals prenatally exposed to alcohol, therefore, it may be that the moderate exposure on G12 used in the current study is not sufficient to increase intake when EtOH is sweetened. Unfortunately, the previous studies using a G17-20 PAE do not report the pregnant dam BECs, so it is difficult to assess whether the exposures were low, moderate, or heavy PAE. However, Chotro and Arias (2003) examined alterations in EtOH intake following different prenatal (G17-G20) doses of EtOH (1g/kg or 2g/kg), and found sex differences as a result of PAE; namely, increased EtOH intake in female offspring was evident following low-dose PAE, whereas male offspring showed an increase when exposed to the higher dose PAE (Chotro and Arias, 2003). They classify this G17-G20 as a moderate exposure, however, conflicting reports equate 1g/kg in a rat to anywhere from 2-4 standard drinks in a human (Abate et al., 2008), which would be more comparable to a 4-day binge-like exposure. Conversely, our model is characterized as a *moderate* exposure, in which rats are exposed one time with a target BEC range between 60-120 mg/dL, much lower in amount and frequency relative to other studies (Valenzuela et al., 2012). Another condition to consider is EtOH concentrations used for assessments of PAE effects on intake. Gore-Langton and Spear (2019) found that PAE (G17-G20) adult males had a higher preference for and intake of 5% than 10% EtOH solution whereas their female counterparts showed less preference and consumed less of 5% EtOH solution. This suggests that PAE animals, in particular males, may preferentially consume solutions containing lower concentrations of EtOH, which could be supported by other studies that also reported increases in EtOH intake of 5-6% EtOH following PAE (Chotro and Arias, 2003; Fabio et al., 2015; Fabio et al., 2013). Interestingly, we found that juvenile PAE groups consumed more *EtOH only* compared to counterparts, which may be an age-dependent effect whereby juveniles have a propensity to consume higher concentrations of EtOH compared to adults (Truxell et al., 2007). Furthermore, it is well-documented in the literature that adolescent animals consume more EtOH than adults, however this adolescent-typical high intake begins to decline around P40 (Doremus et al., 2005; Vetter et al., 2007). Since we began testing juveniles at ∼P25, with the last EtOH intake session occurring ∼P36, and adolescent animals began intake testing P45, it is possible that we may have missed a crucial age range of heightened adolescent EtOH intake in the present study. It is also possible that PAE animals that went through moderate PAE may require more sessions for a significant increase in EtOH intake to occur, as seen with other drinking paradigms (Abate et al., 2008; Chotro and Arias, 2003; Fabio et al., 2015; Fabio et al., 2013; Gore-Langton and Spear, 2019).

In the first intake experiment, we used a 10% EtOH + SS solution that contains sucrose and saccharin, and it is possible that moderate G12 PAE animals may consider this solution less palatable than their air-exposed counterparts. However, intake of ‘supersac’ solution did not differ as a function of prenatal exposure in males and females at any age, suggesting that sweetened solutions were equally palatable for PAE animals and air controls. Given sex differences in preference for various EtOH concentrations in PAE offspring, further studies of PAE effects on EtOH intake should utilize a two- or three-bottle choice paradigm, allowing animals to choose between different concentrations, both sweetened and unsweetened, ultimately providing a more detailed insight into their preferences and behaviors.

An alternative explanation of the decreased intake of sweetened EtOH in PAE adolescents may involve PAE-associated changes in sensitivity to desired effects of EtOH. Indeed, late adolescent animals following a high-dose G12 PAE demonstrated enhanced sensitivity to the socially facilitating and anxiolytic effects of low-dose EtOH (0.5-0.75 g/kg), with these effects evident at BECs between 20 and 45 mg/dL (Mooney and Varlinskaya, 2018). In the present study, intake of sweetened EtOH solution by PAE adolescents resulted in the same BEC range (25-30 mg/dL), suggesting that PAE animals might limit their EtOH intake to achieve desired, presumably anxiolytic effects of EtOH. Clearly, more studies focused on changes in sensitivity to different EtOH effects, including EtOH-induced anxiolysis, following moderate G12 PAE are needed for better understanding of the observed PAE-associated decreases in intake of sweetened EtOH.

Research focused on the development of chronic pain in individuals with FASD suggests that PAE primes the central nervous system to produce aberrant immune responses after a second “hit” to the system, resulting in heightened pain sensitivity (Noor et al., 2017). We initially hypothesized that PAE animals would exhibit heightened tactile sensitivity following postnatal EtOH intake (second “hit”) relative to control animals. While we observed no effect of PAE on mechanical thresholds, similar to previous reports (Noor et al., 2017), we did observe reduced mechanical thresholds after postnatal EtOH experience in all ages. Interestingly, assessments of mechanical thresholds in pre- and postnatal EtOH-naïve animals revealed either a subtle increase in mechanical threshold or stable responding during repeated weekly von Frey tests. Ultimately, these findings suggest that mechanical sensitivity developed as a consequence of post-natal EtOH intake. Furthermore, while air (control) animals consumed more sweetened EtOH than the PAE groups (as discussed above), animals in both prenatal conditions exhibited similar declines in mechanical thresholds, regardless of age. This suggests that despite lower EtOH intake, PAE animals developed mechanical sensitivity at rates comparable to air-exposed controls, suggesting an enhanced sensitivity in response to postnatal EtOH intake in the PAE group. More studies are needed, however, for better understanding of the effects of PAE on sensitivity to EtOH-induced mechanical sensitivity across ontogeny.

There are several limitations of the present study that should be noted. All drinking experiments consisted of six 30-minute sessions that may not be enough to capture PAE-associated increases in EtOH intake. Perhaps more, not necessarily longer, sessions may be needed, since animals tend to consume the most EtOH in the first 30 minutes of most drinking paradigms (Jeanblanc et al., 2019). Furthermore, it is difficult to directly compare the animals’ preference for sweetened vs unsweetened EtOH, given that our drinking paradigm was not conducted in a two-bottle (or more) choice paradigm, instead done with separate cohorts of animals. Despite this, animals drinking either unsweetened EtOH or SS experienced significantly less handling than animals drinking sweetened EtOH solution who were assessed in the LDB or EPM tests of anxiety as well as Von Frey test prior to drinking sessions. A general limitation is the potential floor effect observed in the LDB and EPM data in control animals, as this creates a challenge to detect an increase in anxiety-like behaviors in experimental groups, although we observed both increased and decreased anxiety-like responses respectively in these assays. Furthermore, while we did our best to parse apart the effects of postnatal EtOH intake on mechanical threshold by including EtOH-naïve animals, future studies should assess mechanical thresholds in parallel, where some animals are exposed to a postnatal EtOH intake paradigm and others are exposed to water or a caloric-equivalent solution.

In conclusion, this work contributes to our understanding that even limited and small amounts of PAE may lead to long-lasting neurobehavioral alterations. We have previously found that a single G12 moderate PAE produces a variety of cellular, molecular, and neurophysiological alterations in brain areas involved in anxiety, reward, and pain (basolateral and central amygdala, as well as the prelimbic cortex) in a sex- and age-dependent manner (Przybysz et al., 2023; Rouzer and Diaz, 2022; Winchester and Diaz, 2025), which may contribute to the impairments in behavior observed in this study. Importantly, future research assessing PAE-induced anxiety, substance misuse, and heightened pain sensitivity has the potential to identify underlying mechanisms, ultimately advancing targeted therapeutics for individuals living with FASD.

## Acknowledgements

This work was supported by National Institute on Alcohol Abuse and Alcoholism (NIAAA) R01 AA028566, R01 AA028566-S, P50 AA017823, and F31 AA032198.

