## Supplemental Materials for "Age- and sex- dependent effects of moderate gestational day 12 prenatal alcohol exposure on anxiety-like behaviors, ethanol intake, and mechanical sensitivity"

**Supplementary Table 1**. Elevate plus maze risk assessment behaviors reported as mean (SEM).

|  | **Protected Head**  **Dips** | **Unprotected Head Dips** | **Protected Stretch Attends** | **Unprotected Stretch Attends** |
| --- | --- | --- | --- | --- |
| **Juveniles** | | | | |
| Air Males | 0.875 ± 0.398 | 1.625 ± 0.532 | 2.125 ± 0.667 | 1.750 ± 0.590 |
| PAE Males | 0.429 ± 0.297 | 0.714 ± 0.286 | 1.00 ± 0.436 | 0.750 ± 0.313 |
| Air Females | 1.00 ± 0.327 | 1.429 ± 0.202 | 2.250 ± 0.675 | 0.625 ± 0.324 |
| PAE Females | 1.375 ± 0.625 | 0.875 ± 0.398 | 2.625 ± 0.865 | 0.375 ± 0.183 |
| Exposure  Sex  Exposure x Sex | *F* _1,27_ = 0.007, *p* = 0.936  *F* _1,27_ = 1.479, *p* = 0.234  *F* _1,27_ = 0.869, *p* = 0.359 | *F* _1,26_ = 3.481, *p* = 0.073  *F* _1,26_ = 0.002, *p* = 0.964  *F* _1,26_ = 0.207, *p* = 0.653 | *F* _1,27_ = 0.293, *p* = 0.593  *F* _1,27_ = 1.593, *p* = 0.218  *F* _1,27_ = 1.170, *p* = 0.289 | *F* _1,28_ = 2.672, *p* = 0.113  *F* _1,28_ = 3.847, *p* = 0.060  *F* _1,28_ = 0.962, *p* = 0.335 |
| **Adolescents** | | | | |
| Air Males | 2.50 ± 0.779 | 0.785 ± 0.226 | 3.750 ± 0.726 | 0.750 ± 0.250 |
| PAE Males | 2.875 ± 1.141 | 2.50 ± 0.779 | 3.875 ± 1.043 | 1.625 ± 0.596 |
| Air Females | 2.625 ± 0.532 | 2.250 ± 0.818 | 4.750 ± 0.840 | 2.00 ± 0.598 |
| PAE Females | 3.00 ± 0.779 | 3.875 ± 1.109 | 4.875 ± 0.789 | 3.00 ± 0.500 |
| Exposure  Sex  Exposure x Sex | *F* _1,28_ = 0.201, *p* = 0.657  *F* _1,28_ = 0.022, *p* = 0.882  *F* _1,28_ = 0.000, *p* > 0.999 | ***F* _1,28_ = 4.129, *p* =0.052**  *F* _1,28_ = 2.956, *p* = 0.096  *F* _1,28_ = 0.000, *p* > 0.999 | *F* _1,28_ = 0.021, *p* = 0.886  *F* _1,28_ = 1.360, *p* = 0.254  *F* _1,28_ = 0.000, *p* =0.999 | *F* _1,28_ = 3.431, *p* = 0.075  ***F* _1,28_ = 6.725, *p* = 0.015**  *F* _1,28_ = 0.015, *p* = 0.903 |
| **Adults** | | | | |
| Air Males | 2.571 ± 0.429 | 2.125 ± 0.667 | 4.875 ± 0.581 | 0.875 ± 0.295 |
| PAE Males | 2.375 ± 0.532 | 1.625 ± 0.420 | 4.00 ± 0.535 | 1.125 ± 0.350 |
| Air Females | 2.250 ± 0.648 | 2.00 ± 0.802 | 4.875 ± 0.990 | 1.125 ± 0.398 |
| PAE Females | 3.250 ± 0.491 | 2.750 ± 0.996 | 5.750 ± 0.818 | 2.625 ± 0.755 |
| Exposure  Sex  Exposure x Sex | *F* _1,27_ = 0.556, *p* = 0.463  *F* _1,27_ = 0.264, *p* = 0.612  *F* _1,27_ = 1.232, *p* = 0.277 | *F* _1,28_ = 0.028, *p* = 0.869  *F* _1,28_ = 0.444, *p* = 0.512  *F* _1,28_ = 0.693, *p* = 0.412 | *F* _1,27_ = 0.000, *p* = 0.999  *F* _1,27_ = 1.348, *p* = 0.256  *F* _1,27_ = 1.348, *p* = 0.256 | *F* _1,28_ = 3.267, *p* = 0.082  *F* _1,28_ = 3.267, *p* = 0.082  *F* _1,28_ = 1.667, *p* = 0.207 |

Significant effects are presented in **bold.**

**Supplementary Table 2.** Relationship between ethanol + supersac intake and blood ethanol concentrations

| Age | Male | | Female | |
| --- | --- | --- | --- | --- |
|  | Air | PAE | Air | PAE |
| Juvenile | **R = 0.498**  **R^2^ = 0.248**  ***F* _1,14_ = 4.611**  ***p* = 0.0498** | R = 0.7236  R^2^ = 0.0092  *F* _1,14_ = 0.1302  *p* = 0.7236 | **R = 0.656**  **R^2^ = 0.430**  ***F* _1,14_ = 10.56**  ***p* = 0.0058** | **R = 0.923**  **R^2^ = 0.853**  ***F* _1,14_ = 81.03**  ***p* < 0.0001** |
| Adolescent | R = 0.4473  R^2^ = 0.200  *F* _1,14_ = 3.501  *p* = 0.0824 | **R = 0.7462**  **R^2^ = 0.5569**  ***F* _1,14_ = 17.59**  ***p* = 0.0009** | **R = 0.8791**  **R^2^ = 0.7727**  ***F* _1,14_ = 47.61**  ***p*** **< 0.0001** | R = 0.3549  R^2^ = 0.1258  *F* _1,14_ = 2.015  *p* = 0.1776 |
| Adult | **R = 0.5200**  **R^2^ = 0.2704**  ***F* _1,14_ = 5.188**  ***p* = 0.0390** | **R = 0.5832**  **R^2^ = 0.3402**  ***F* _1,14_ = 7.217**  ***p* = 0.0177** | **R = 0.6259**  **R^2^ = 0.3918**  ***F* _1,14_ = 9.017**  ***p* = 0.0095** | **R = 0.5792**  **R^2^ = 0.3355**  ***F* _1,14_ = 7.067**  ***p* = 0.0187** |

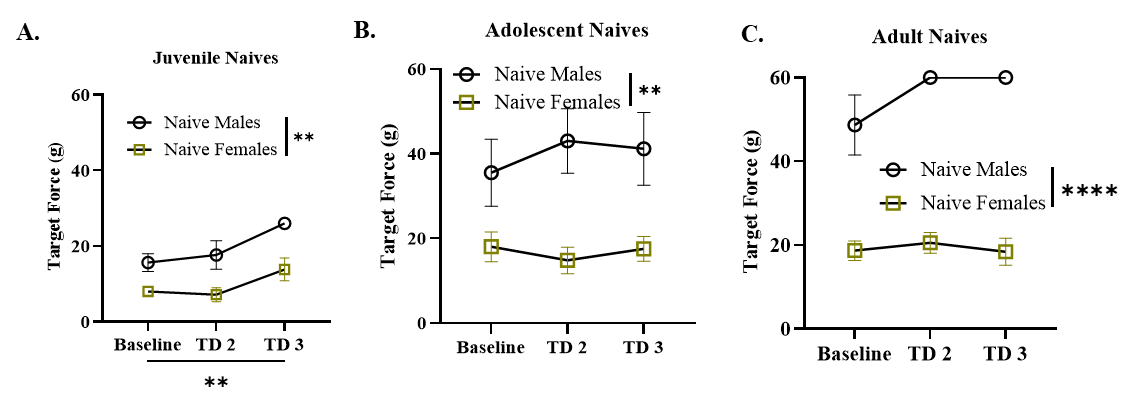

**Supplementary Figure 1: Von Frey – Ethanol naïve**. **(A-C)** A main effect of sex was observed in juveniles, adolescents, and adult naïve animals, where males had a higher mechanical threshold compared to females. **(A)** An effect of test day was revealed in juveniles, where threshold increased over time. * denoted p ≤ 0.05. Bars represent standard error of the mean. PAE = Prenatal alcohol exposure.
